# DrugDomain 2.0: comprehensive database of protein domains-ligands/drugs interactions across the whole Protein Data Bank

**DOI:** 10.1101/2025.07.03.663025

**Authors:** Kirill E. Medvedev, R. Dustin Schaeffer, Nick V. Grishin

## Abstract

Proteins carry out essential cellular functions – signaling, metabolism, transport – through the specific interaction of small molecules and drugs within their three-dimensional structural domains. Protein domains are conserved folding units that, when combined, drive evolutionary progress. The Evolutionary Classification Of protein Domains (ECOD) places domains into a hierarchy explicitly built around distant evolutionary relationships, enabling the detection of remote homologs across the proteomes. Yet no single resource has systematically mapped domain-ligand interactions at the structural level. To fill this gap, we introduce DrugDomain v2.0, an updated comprehensive resource, that extends earlier releases by linking evolutionary domain classifications (ECOD) to ligand binding events across the entire Protein Data Bank. We also leverage AI-driven predictions from AlphaFold to extend domain-ligand annotations to human drug targets lacking experimental structures. DrugDomain v2.0 catalogs interactions with over 37,000 PDB ligands and 7,560 DrugBank molecules, integrates more than 6,000 small–molecule–associated post-translational modifications, and provides context for 14,000+ PTM-modified human protein models featuring docked ligands. The database encompasses 43,023 unique UniProt accessions and 174,545 PDB structures. The DrugDomain data is available online: https://drugdomain.cs.ucf.edu/ and https://github.com/kirmedvedev/DrugDomain.

## 1. Introduction

Studying how small molecules and drugs interact with protein structural domains lies at the heart of understanding both molecular function and guiding drug discovery. Through the binding of endogenous cofactors, metabolites, or exogenous drugs within their structural three-dimensional domains, proteins participate in a variety of vital cellular processes, including signaling, metabolism, and transport. Protein domains are conserved structural, functional, and evolutionary units that serve as the essential building blocks for protein diversity and adaptation [1]. The different ways in which protein domains can be combined provide a powerful mechanism for evolving new protein functions and shaping cellular processes [2]. Identifying and categorizing protein domains based on their evolutionary relationships can enhance our understanding of protein function. This is achieved by examining the established functions of their homologs. Until recently, major structure-based classifications of protein domains were primarily centered on categorizing experimentally determined protein structures, e.g., SCOP [3] and CATH [4]. Our team has developed and maintains the Evolutionary Classification of Protein Domains database (ECOD), whose key feature is its emphasis on distant homology, which culminates in a comprehensive database of evolutionary relationships among categorized domains’ topologies [5, 6]. Mapping the protein-ligand interactions at the domain level can reveal the mechanistic basis of protein function and inform structure-based drug discovery.

Artificial intelligence provides powerful tools for scientific research across diverse fields, and structural computational biology is no exception. AlphaFold (AF) has revolutionized structural biology by demonstrating atomic-level precision in protein structure prediction and becoming an indispensable tool in the field [7]. Leveraging AF models, ECOD stands out as one of the first databases to provide comprehensive domain classifications for both the entire human proteome [8] and the complete proteomes of 48 additional model organisms [6]. Recently, The Encyclopedia of Domains (TED) [9] was released - a comprehensive resource for the identification and classification of protein domains within the AlphaFold Database [10]. This advancement by AlphaFold has significantly broadened the scope of computational structural biology, enabling diverse applications such as drug discovery, drug target prediction, and the analysis of protein-protein and protein-ligand interactions [11, 12]. The new release of AlphaFold3 has further improved the accuracy of protein structure and protein-ligand interaction predictions [7].

As of today, no available resource reports interactions between protein structural domains (based on evolutionary classification) and ligands. With the latest advances in AI-based methods for predicting protein structure and protein-ligand interactions, we are witnessing a paradigm shift where computational approaches achieve performance levels nearly comparable to those of experimental methods. Here we present DrugDomain v2.0 (https://drugdomain.cs.ucf.edu/), a comprehensive database detailing the interactions of structural protein domains with a wide array of small organic (including drugs) and inorganic compounds, and – unlike previous versions – covering the full breadth of the Protein Data Bank. Our dataset encompasses all ligands in the Protein Data Bank that interact with protein structures. The database also provides domain-drug interactions for AlphaFold models of human drug targets without solved experimental structures [13]. It also features over 6,000 small-molecule binding-associated PTMs and more than 14,000 PTM-modified human protein models with docked ligands [14]. In total, the database now encompasses 43,023 unique UniProt accessions, 174,545 PDB structures, 37,367 PDB ligands, and 7,561 DrugBank molecules. We believe this resource can serve as a foundation for a range of forward-looking studies – including drug repurposing, the development of improved docking protocols, and the analysis of post-translational modifications in protein-ligand interactions.

## 2. Materials and Methods

### 2.1. Data collection and analysis

The comprehensive list of ligands and small molecule components found in Protein Data Bank [15] was retrieved from Chemical Component Dictionary [16]. All PDB entries containing these ligands and small molecules’ InChI Key and SMILES formulas were obtained using rcsb-api [17]. Using InChI Keys and SMILES, we retrieved accession numbers for each small molecule from the following databases, where available: DrugBank [18], PubChem [19], ChEMBL [20]. In the DrugDomain database, we use the PDB ligand ID as a primary identifier for the small molecule (for example, NAD, 2I4, etc.). Alternatively, we use DrugBank accession for cases when the PDB ligand ID is unknown. Additionally, drug action data were retrieved from DrugBank and affinity data from BindingDB [21]. Chemical classification of small-molecule components was obtained from the ClassyFire database [22] and includes the four top levels of the classification: kingdom, superclass, class and subclass. 2D diagrams of ligand-protein interactions (LigPlots) were generated using LigPlot+ as in v1.0 and v.1.1 [23].

For each ligand-protein (PDB structure) pair, residues located within 5 Å of the atoms of the small molecule were identified using BioPython [24]. Interacting residues were mapped to structural domains from ECOD database v292 (08302024) [5] and reported in DrugDomain. For ligand– protein pairs lacking experimentally determined structures, we used AlphaFold models and the AlphaFill algorithm [25] to transplant missing ligands from PDB structures into these models based on sequence and structural similarity. This process was performed in DrugDomain v1.0 for the subset of human proteins known to interact with small molecules and drugs from DrugBank. The methodology and implementation of this approach into the DrugDomain database was described previously [13]. To calculate ligand-interacting statistics based on the number of domains, we counted the UniProt-accessioned proteins that included a specific number of ECOD domains interacting with the ligand.

In DrugDomain v1.1 we explored the effect of post-translational modifications (PTMs) on small molecule binding for the subset of human proteins from v1.0. We used recent AI-based approaches for protein structure prediction (AlphaFold3 [7], RoseTTAFold All-Atom [26], Chai-1 [27]) and generated 14,178 models of PTM-modified human proteins with docked ligands [14]. To do that, we identified PTMs within 10 Å of all atoms of each small molecule bound to human proteins in the subset of human proteins from v1.0. The overall number of identified small molecule binding-associated PTMs was 6,131. Overall, we generated 1,041 AlphaFold3, 9,169 RoseTTAFold All-Atom and 3,968 Chai-1 PTM-modified models. Each DrugDomain webpage includes a placeholder indicating the availability of PTM data for each protein–small molecule combination presented in the DrugDomain database. If PTM data is available, there is a link “List of drug binding-associated PTMs”; otherwise, it states “No PTM data available”. The major novelty of DrugDomain v2.0, compared to previous versions (v1.0 and v1.1), is the inclusion of domain– ligand interaction data across the entire Protein Data Bank. In addition to the human protein subset and small molecules from DrugBank (v1.0 and v1.1), we incorporated all ligands from the PDB and all experimental protein structures that interact with these ligands.

## 3. Results and Discussion

### 3.1. DrugDomain v2.0 statistics and features

DrugDomain v2.0 includes the following major types of data related to interactions between protein domains and small molecule components. First, the new version of DrugDomain reports domain-ligand interactions for all PDB entries containing ligand entities, including both organic small molecules and inorganic components. Thus, we expanded the scope of the database to encompass not only protein-drug interactions but also interactions between protein domains and all ligand entities that are present in PDB. Second, the v2.0 reports domain-drug interactions for AlphaFold models of human drug target proteins lacking experimentally determined structures [13]. Third, it includes over 6,000 small molecule binding-associated PTMs identified in the human proteome and over 14,000 PTM-modified human proteins with docked ligands generated using recent AI-based approaches (AlphaFold3 [7], RoseTTAFold All-Atom [26], Chai-1 [27]) [14]. To help users navigate between different types of data, we created a detailed tutorial (https://github.com/kirmedvedev/DrugDomain/wiki/DrugDomain-database-Tutorial).

DrugDomain database v2.0, includes 43,023 unique UniProt accessions [28], 174,545 PDB structures (over 70% of all experimental protein structures), 37,367 ligands from PDB, 7,561 DrugBank molecules (over 50% of all small molecule drugs in DrugBank) (Fig. 1).

**Figure 1.**
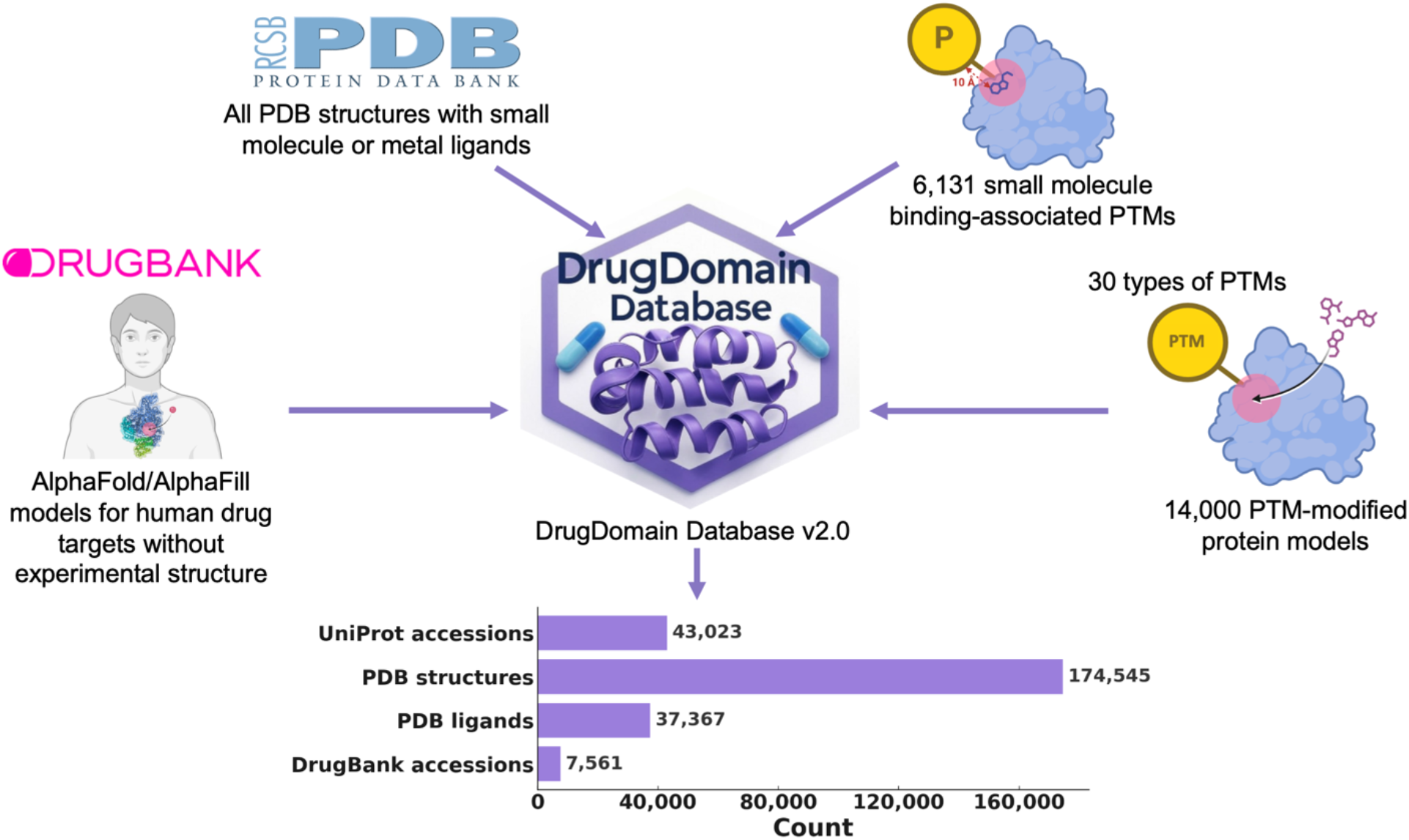
DrugDomain database v2.0 data types and statistics.

DrugDomain includes two types of hierarchy: protein and molecule-centric. The complete lists of proteins and small molecules can be accessed through the top menu. There are two types of molecule lists – by DrugBank accession and by PDB ligand ID. The protein or molecule can be searched using the search field on the main page or the quick search option at the navigation bar. The search can be conducted using UniProt (e.g. Q03181), PDB ligand (e.g. ATP), DrugBank accessions (e.g. DB00171), or SMILES formula. The search by UniProt accession returns a list of ligands known or predicted to interact with the query protein, along with key data for each ligand: PDB ID; DrugBank, PubChem, and ChEMBL accessions; molecule name; drug action; and affinity. The molecule search (by PDB ligand ID, DrugBank accession, or SMILES formula) returns a list of proteins known or predicted to bind the query molecule, along with key data for ligand and protein. Both search types return links to DrugDomain data pages, which provide key ligand information, including its chemical classification, and list PDB structures and/or AlphaFold models known or predicted to bind the ligand. The list of the structures includes PDB/AF accession, downloadable PyMOL [29] script, which shows ECOD domains and residues interacting with the ligands; a list of ECOD domains interacting with the molecule with links to the ECOD database, names of corresponding ECOD X-groups (possible homology level) and 2D diagrams of ligand–protein interactions (LigPlots). DrugDomain data webpage also includes a link to a list of drug-binding-associated post-translational modifications (PTMs) where available [14]. This list contains information about each PTM and links to PyMOL sessions with models of modified proteins generated by AlphaFold3, RoseTTAFold All-Atom or Chai-1. PyMOL sessions include PTM-modified residues, the ligand, and mapped ECOD domains, each shown in different colors.

The taxonomic distribution of proteins reported in the DrugDomain database v2.0 revealed the prevalence of eukaryotic and bacterial proteins (Fig. 2A). *Pseudomonadota* or proteobacteria are one of the most abundant phyla of Gram-negative bacteria, which are naturally found as pathogenic and free-living genera [30]. Thus, proteins from these bacteria are important targets for antibacterial therapy against human pathogens, and PDB entries of these proteins bound to various antibiotics comprise a significant fraction of the Protein Data Bank. Bacteria belonging to the phylum *Bacillota* can make up 11-95% of the human gut microbiome [31] and play key roles in energy extraction. They have also been associated with the development of diabetes and obesity [32], making them potential therapeutic targets. Finally, the third-largest phylum in terms of the number of PDB structures with ligands is *Actinomycetota* (or Actinobacteria). These bacteria are major contributors to the biological buffering of soils and the source of many antibiotics [33]. Similarly, there are three largest eukaryotic phyla: Chordata includes humans and various model organisms such as mice and rats; Ascomycota is the largest phylum of fungi, which are the source of antibiotics like penicillin, and particular species are used to produce immunosuppressants and other medicinal compounds [34]; *Streptophyta* phylum includes green algae and the land plants. The distribution of ECOD domains from experimental structures interacting with ligands is shown in Figure 2B. The top three largest ECOD A-groups include α/β three-layered sandwiches, α+β two layers and α+β complex topology. The α/β three-layered sandwich architecture is represented mainly by Rossmann-like proteins. In our earlier work, we showed that these proteins perform diverse functions and interact with most superclasses of organic molecules [35, 36]. Most small molecules that interact with domains of the α+β complex topology target protein kinases, which are among the most druggable proteins in the human proteome; therefore, their structures are abundant in the Protein Data Bank [37, 38]. The α+β two-layer architecture includes heat shock proteins (HSP), which play a critical role as molecular chaperones and are important targets for anticancer chemotherapy [39].

**Figure 2.**
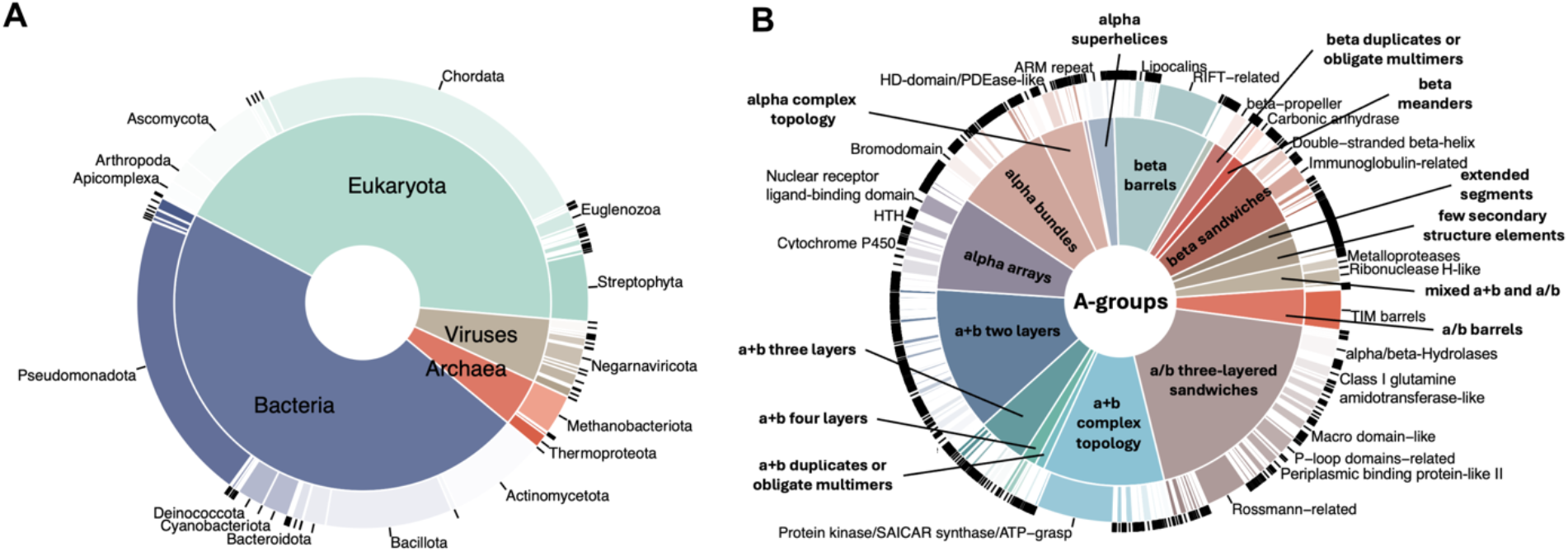
DrugDomain v2.0 statistics. **(A)** Taxonomic distribution of proteins reported in the DrugDomain database, by UniProt population. The inside pie shows the distribution of superkingdoms, and the outside donut shows the distribution of phyla. **(B)** Distribution of ECOD domains from experimentally determined PDB structures, interacting with ligand, stratified by architecture (inside pie) and homologous group (outside donut).

Analysis of domains from experimentally determined PDB structures and the ClassyFire superclasses of the organic compounds they interact with revealed the three most common superclasses [22] in Protein Data Bank: Organoheterocyclic compounds, Organic oxygen compounds, Organic acids and derivatives (Fig. 3). The largest fraction of domains interacting with compounds from the majority of superclasses belongs to α/β three-layered sandwiches, α+β two layers and α+β complex topology ECOD architecture types, which were discussed above. The superclass Organoheterocyclic compounds includes atorvastatin, a lipid-lowering drug that reduces the risk of myocardial infarction, stroke, and other cardiovascular diseases [40].

**Figure 3.**
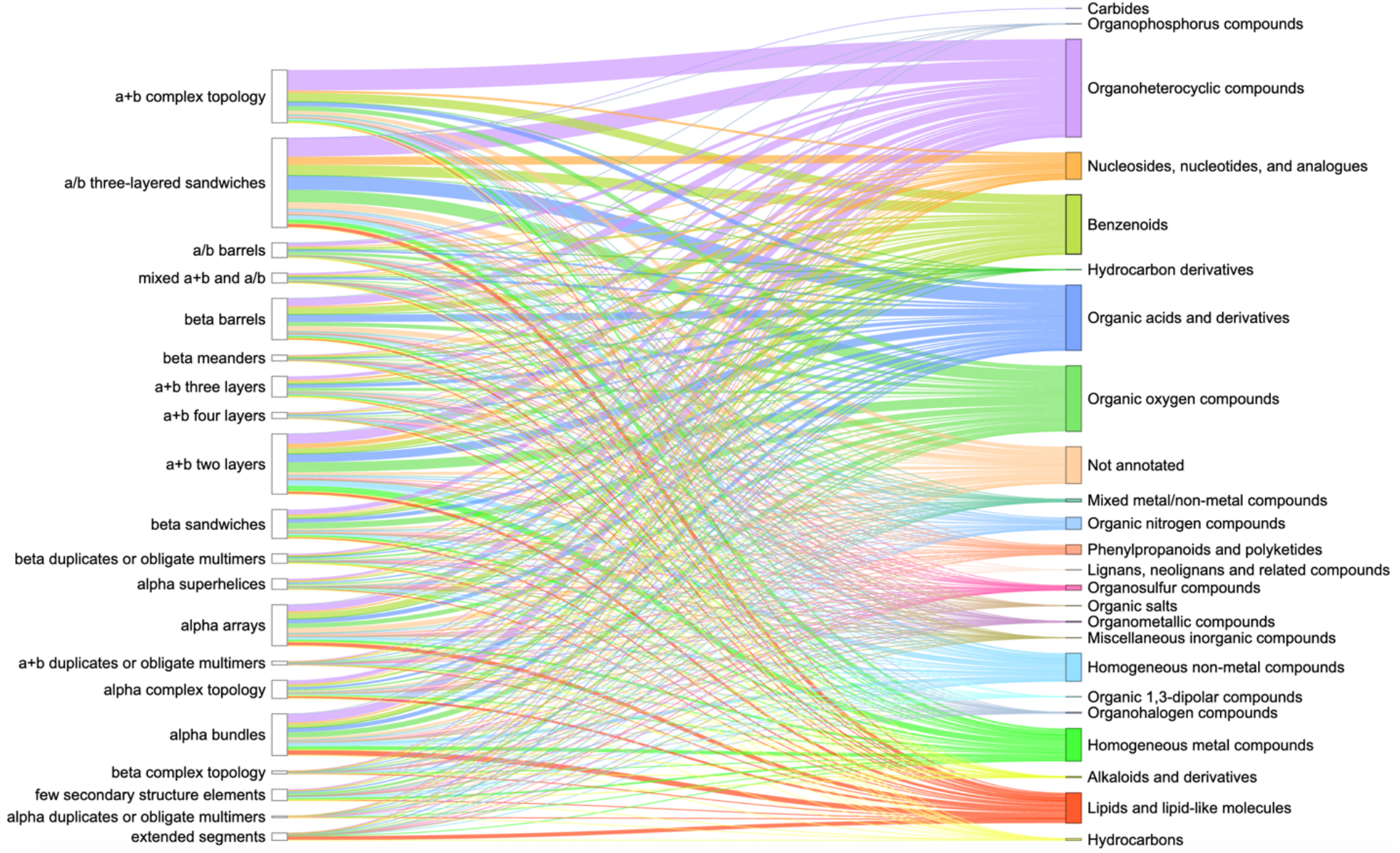
ECOD A-groups (left column) of experimental PDB structures and superclasses of organic molecules according to ClassyFire classification (right column). Each superclass and the lines pointed toward it are denoted by separate color. The thickness of the lines shows the number of PDB ligands interacting with domains from ECOD A-groups.

Erythromycin is a broad-spectrum antibiotic in the Organic oxygen compounds superclass and is widely used to treat infections caused by both Gram-positive and Gram-negative bacteria [41]. Finally, Arbaclofen – a member of the Organic acids and derivatives superclass - is a drug that is used in the treatment of autism [42].

### 3.2. Number of domains mediating ligand interactions in Protein Data Bank

Protein domains are conserved structural units that serve as the fundamental evolutionary and architectural building blocks of proteins. Understanding how ligands bind – specifically, which domains are involved and how many mediate the interaction – is crucial for uncovering protein function and guiding drug discovery. Overall ligand-interacting statistics were calculated for each protein, based on the number of interacting ECOD domains associated with its UniProt accession (Fig. 4). Our results revealed that the majority of proteins with assigned ECOD domains bound ligands using one or two domains. Our observation is consistent with previous research, which indicates that most drug targets bind via a limited set of prevalent domains [43]. Moreover, it is noteworthy that, under the ECOD classification, protein kinases – the most druggable targets in the human proteome – are characterized by a single structural domain [38]. This contributes to their significant representation among proteins with one ligand-interacting domain. In contrast, other structural classifications divide these proteins into two domains [4]. It is important to note, however, that experimentally determined PDB structures may not always accurately reflect ligand coordination, as only a part of the protein is often included in the experimental structure.

**Figure 4.**
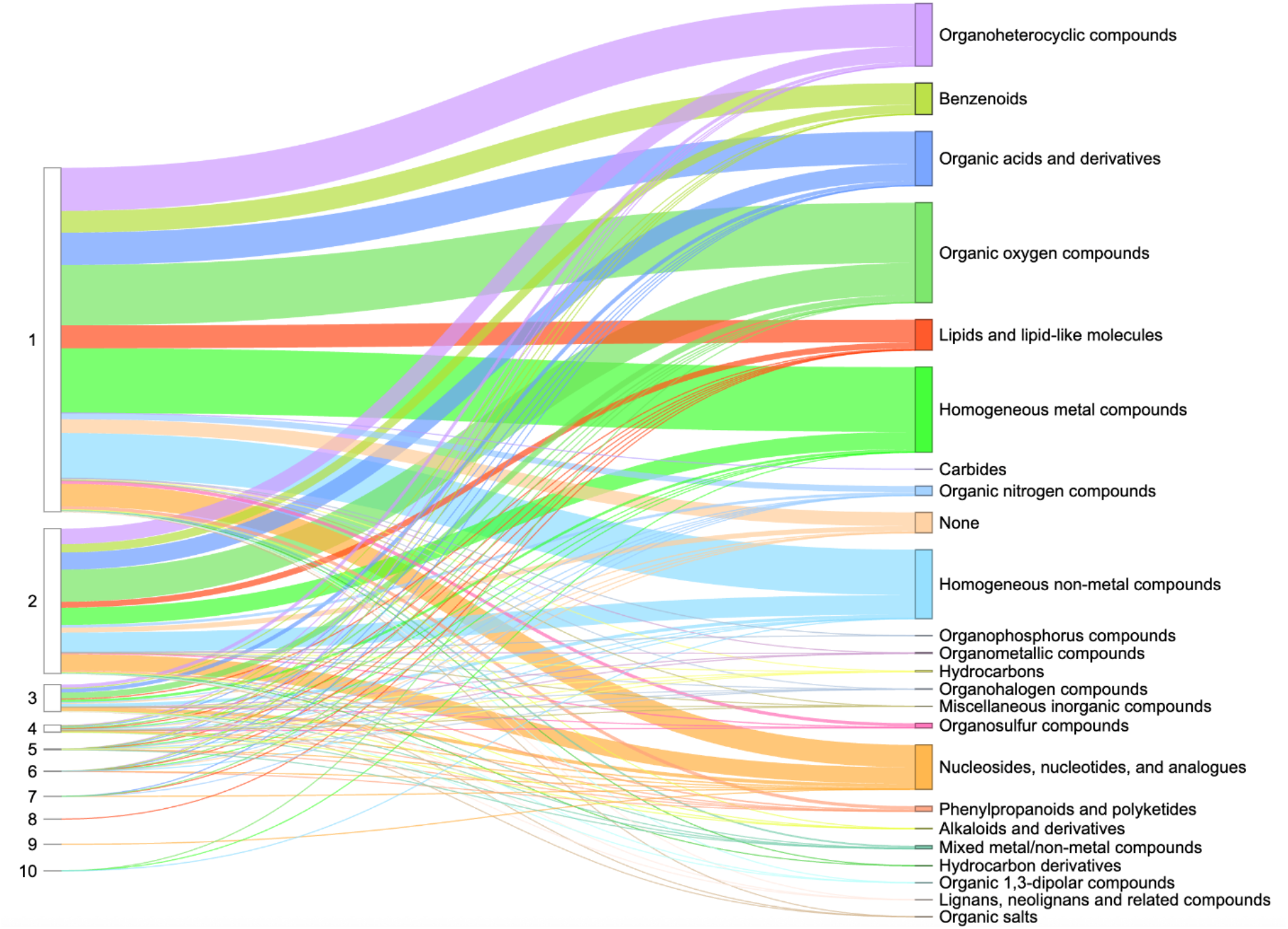
Ligand-interacting statistics by number of domains per UniProt accession in Protein Data Bank. The left column shows the number of ligand-interacting domains, the right column shows the superclasses of organic molecules according to ClassyFire classification. The thickness of the lines indicates the number of UniProt accessions.

Our analysis of ligand-interacting statistics indicated that proteins deposited in the Protein Data Bank contain a range of one to ten ECOD domains involved in ligand interaction (Fig. 4). Such a large number of interacting domains (ten) can bind a single ligand when the protein forms a channel or pore structure. For example, human mitochondrial RNA splicing 2 (Mrs2) channel (Fig. 5A-C) enables Mg^2+^ permeation across the inner mitochondrial membrane and is crucial for mitochondrial metabolic function [44], illustrating how a channel structure can accommodate interactions with multiple domains. Dysregulated Mg^2+^ levels in humans are implicated in various diseases [45], as mitochondria are the primary site of ATP production in eukaryotic cells – a process critically dependent on Mg^2+^ as a cofactor. The cation also commonly forms complexes with cellular nucleotides [46]. Mrs2 exists as homopentamers, with each monomer featuring two C-terminal transmembrane helices [46]. Structurally, each monomer contains two ECOD domains: an N-terminal “CorA soluble domain-like” domain and a C-terminal transmembrane domain (Fig. 5C). Mg^2+^ is coordinated near the borders of two domains of each monomer and interacts with each domain of homopentamer (Fig. 5B).

**Figure 5.**
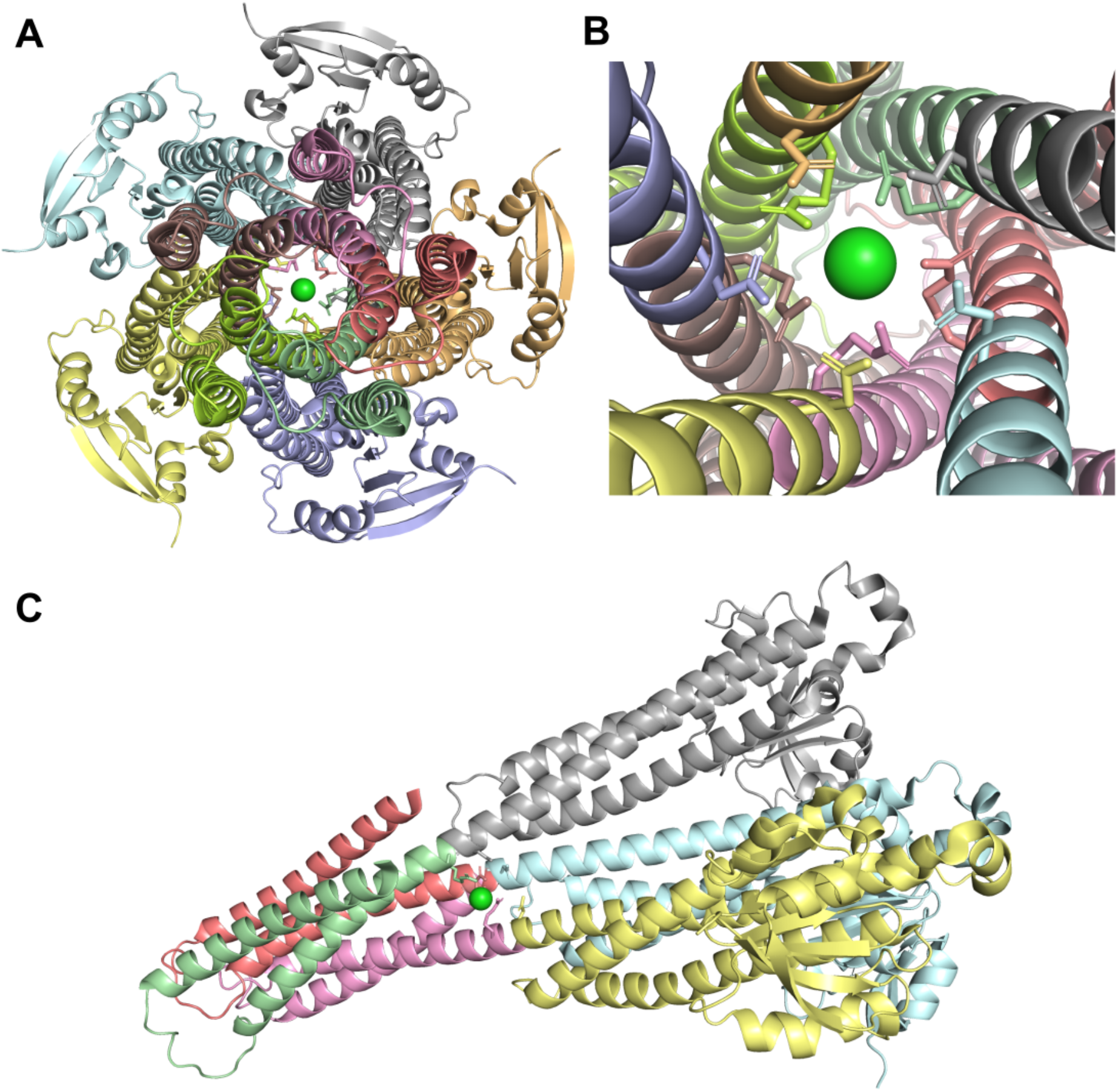
Structure of the human mitochondrial Mrs2 channel (PDB: 8IP5). **(A)** Channel view of Mrs2 with protein colored by ECOD domains, Mg^2+^ ion is shown in green, and sticks show interacting residues. **(B)** Close-up channel view of Mrs2. **(C)** Side view of Mrs2 showing three out of five monomers. Chains C, D, and E are colored by ECOD domains “

The DrugDomain database allows users to explore all known interactions of a given ligand with all known targets. For example, ATP – one of the most prevalent biological ligands – interacts with 1,035 proteins (https://drugdomain.cs.ucf.edu/molecules/pdb/ATP.html - counted by UniProt accession) and may be coordinated by structurally unrelated (non-homologous) domains (Fig. 6A-D). For example, ubiquitin-like modifier-activating enzyme Atg7 activates two ubiquitin-like proteins, Atg8 and Atg12, and plays a crucial role in autophagy [47]. Figure 6A shows Atg7 (orange), represented by a domain from the Rossmann-related ECOD H-group, bound to Atg8 (blue), represented by the Ubiquitin-Related H-group (beta-Grasp X-group). Atg7 takes part in adenylation of the C-terminal Gly residue of ubiquitin-like proteins, and this step consumes ATP [47]. In Fig. 6B, Cobalamin adenosyltransferase (ATR) is shown as two chains (orange and brown). Each chain comprises a single, almost exclusively α-helical domain that belongs to the Cobalamin adenosyltransferase H-group. ATR catalyzes the adenosylation of cob(I)alamin by ATP, which leads to cobalt−carbon bond formation and the synthesis of coenzyme B_12_ [48]. In Fig. 6C cytoplasmic part of ATP-binding cassette transporter ABCG2 is shown. ABCG2 is a transporter localized to the plasma membrane of cells across multiple tissues and physiological barriers. It mediates translocation of endogenous substrates, modulates the pharmacokinetics of numerous therapeutics, and confers protection against a wide spectrum of xenobiotics, including anticancer drugs [49]. This process is powered by ATP. ATP-binding domains of this protein (grey and cyan) belong to P-loop domains-related H-group and contain the canonical P-loop sequence motif that coordinates ATP molecule. Finally, Fig. 6D shows the ATP phosphoribosyltransferase that forms a homodimer of domains belonging to the Periplasmic binding protein-like II H-group. This protein catalyses the first step of histidine biosynthesis in plants and microorganisms. This is an energetically expensive process requiring 41 ATP equivalents for the synthesis of one histidine molecule [50].

**Figure 6.**
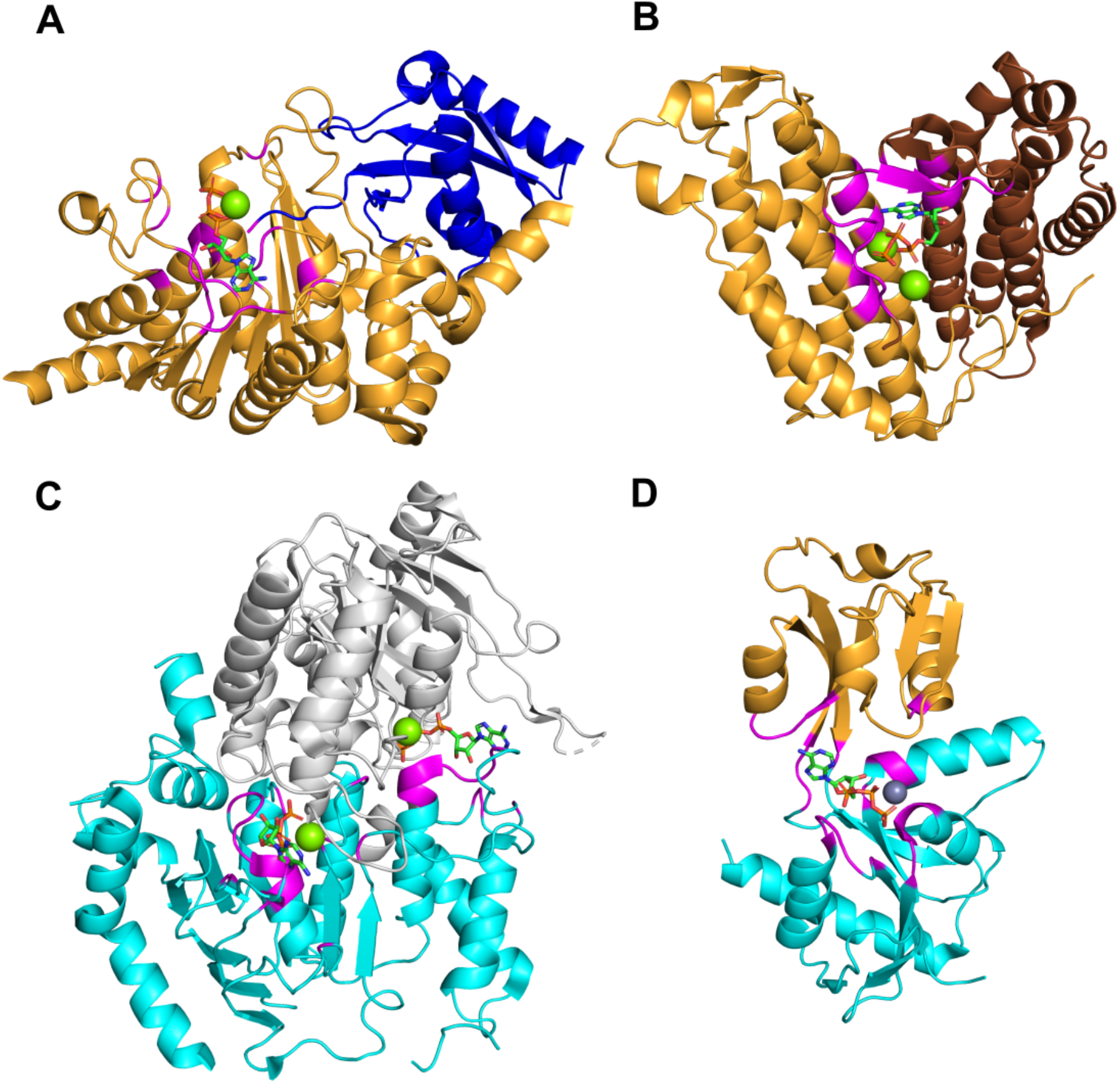
Examples of ATP binding to different proteins. **(A)** Ubiquitin-like modifier-activating enzyme Atg7 bound to Atg8 (PDB: 3VH4). **(B)** Cobalamin adenosyltransferase MMAB (PDB: 6D5K). **(C)** ATP-binding cassette transporter ABCG2 (PDB: 6HZM) **(D)** ATP phosphoribosyltransferase (PDB: 5UBH). All proteins are colored by their ECOD domains. ATP is depicted with sticks and colored by its constituent elements. Residues interacting with ATP are colored in magenta.

## Conclusions

The DrugDomain database version 2.0 represents a comprehensive resource depicting interactions between structural protein domains and small organic (including drugs) and inorganic molecules, and – unlike previous versions – covers the entire Protein Data Bank. It also reports domain-drug interactions for AlphaFold models of human drug targets lacking experimental structures. Additionally, it features over 6,000 small-molecule binding-associated PTMs and more than 14,000 PTM-modified human protein models with docked ligands, generated by state-of-the-art AI-based approaches. DrugDomain database v2.0 includes 43,023 unique UniProt accessions (more than 16-fold increase relative to v1.0), 174,545 PDB structures, 37,367 ligands from PDB, and 7,561 DrugBank molecules. Within experimental PDB structures, the distribution of ECOD domains interacting with ligands was analyzed. This analysis revealed that the top three ECOD A-groups, ranked by the number of ligand-interacting domains, are predominantly α/β three-layered sandwiches (Rossmann fold), α+β two layers (heat shock proteins), and α+β complex topology (kinases). The distribution of domains in experimental PDB structures and their interacting compound superclasses identified the top three categories as Organoheterocyclic compounds, Organic oxygen compounds, and Organic acids and derivatives. Our analysis showed that proteins in the Protein Data Bank exhibit a range of one to ten ECOD domains involved in ligand interaction. All data and protein models are available for viewing and downloading in the DrugDomain database (https://drugdomain.cs.ucf.edu/) and GitHub (https://github.com/kirmedvedev/DrugDomain).

## Competing interests

The authors declare that there are no competing interests associated with the manuscript.

## Acknowledgements

The authors acknowledge the Texas Advanced Computing Center (TACC) at The University of Texas at Austin (https://tacc.utexas.edu/) for providing computational resources that have contributed to the research results reported within this paper.

## Funding

The study is supported by The University of Central Florida College of Engineering and Computer Science (to K.E.M.), grants from the National Institute of General Medical Sciences of the National Institutes of Health GM127390 (to N.V.G.), GM147367 (to R.D.S), the Welch Foundation I-1505 (to N.V.G.), the National Science Foundation DBI 2224128 (to N.V.G.).

## CRediT Author Contribution

**Kirill E. Medvedev:** Conceptualization, Methodology, Software, Validation, Formal analysis, Investigation, Data Curation, Visualization, Writing - Original Draft, Writing - Review & Editing, Project administration, Funding acquisition. **R. Dustin Schaeffer**: Writing - Review & Editing, Funding acquisition. **Nick V. Grishin:** Resources, Funding acquisition.

## Notes

https://drugdomain.cs.ucf.edu/

### Competing Interest Statement

The authors have declared no competing interest.

### Summary of Updates

Methods section expanded; Figures 2,3,5,6 revised; website of the database updated

